# SUMO/deSUMOylation of the BRI1 brassinosteroid receptor modulates plant growth responses to temperature

**DOI:** 10.1101/2022.01.17.476605

**Authors:** Maria Naranjo-Arcos, Moumita Srivastava, Mansi Bhardwarj, Ari Sadanandom, Grégory Vert

## Abstract

Brassinosteroids (BRs) are a class of steroid molecules perceived at the cell surface that act as plant hormones. The BR receptor BRI1 offers a model to understand receptor-mediated signaling in plants and the role of post-translational modifications. Here we identify SUMOylation as a new modification, targeting BRI1 to regulate its activity. BRI1 is SUMOylated *in planta* on two lysine residues and the levels of BRI1-SUMO conjugates are controlled by the Desi3a SUMO protease. We demonstrate that BRI1 is deSUMOylated at elevated temperature by Desi3a, leading to increased BRI1 interaction with the negative regulator of BR signaling BIK1 and enhancing BRI1 endocytosis. Loss of Desi3a or BIK1 results in increased response to temperature elevation, indicating that BRI1 deSUMOylation acts as a safety mechanism necessary to keep temperature responses in check. Altogether, our work establishes BRI1 deSUMOylation as a molecular crosstalk mechanism between temperature and BR signaling, allowing plants to translate environmental inputs into growth response.

**SIGNIFICANCE STATEMENT:** The brassinosteroid (BR) receptor BRI1 provides a paradigm for understanding receptor-mediated signaling in plants and contribution of post-translational modifications. Here, we show that BRI carries SUMO modifications in planta on two intracellular lysine residues and that temperature elevation triggers BRI1 deSUMOylation mediated by the Desi3a SUMO protease. Importantly, BRI1 deSUMOylation leads to downregulation of BR signaling via increased BRI1 interaction with the BIK1 negative regulator and increased BRI1 endocytosis. Loss of BRI1 deSUMOylation in *desi3a* mutants boosts plant responses to heat, indicating that BRI1 deSUMOylation acts as a brake to keep temperature responses in check. Our study uncovers a new post-translational modification targeting BRI1 and sheds light on its functional outcome for environmentally-controlled plant growth.

## INTRODUCTION

Brassinosteroids (BRs) are a class of plant hormones controlling various aspects of plant development and stress responses (1). Genetic, biochemical, and structural biology studies have revealed that BRs are perceived at the cell surface by the BRASSINOSTEROID INSENSITIVE1 (BRI1) leucine-rich-repeat receptor-like kinase (LRR-RLK) (2–7). BR binding to BRI1 promotes its heterodimerization with the LRR-RLK BRI1-ASSOCIATED RECEPTOR KINASE (BAK1) (8, 9) to form a competent receptor complex. *cis* and *trans* phosphorylation of BRI1 and BAK1 fully activates the receptor complex and initiates a protein phosphorylation-mediated signaling cascade, which ultimately regulates the activity of the BRASSINAZOLE RESISTANT(BZR)-family of transcription factors (10, 11). Among these, BZR1 and BES1 were shown to control the expression of thousands of BR-responsive genes important for plant growth and stress response (12, 13).

Inability to properly produce, sense or transduce the BR signal results in characteristic BR-deficient/insensitive phenotypes that include short hypocotyls in the dark, dwarf stature in the light, altered vascular development, prolonged vegetative phase, and reduced male fertility (5, 14–16). In contrast, BR overproduction or enhanced signaling activities are associated with increased growth (7). The precise control of BR perception at the cell surface is therefore crucial to ensure proper plant development and completion of the plant life cycle in an ever-changing environment. As the major BR receptor and a long-lived protein, BRI1 is regulated by several mechanisms to keep its basal activity in check and to desensitize BRI1 after BR signaling. The BRI1 kinase is first kept in its basal state by an autoinhibitory C-terminal tail (17). Phosphorylation of the C-terminal tail upon BR binding likely releases autoinhibition for the full activation of BRI1. BRI1 also interacts with an inhibitory protein named BRI1 KINASE INHIBITOR1 (BKI1) that prevents interaction between BRI1 and BAK1 in resting cells (18). BR binding to BRI1 triggers BKI1 tyrosine phosphorylation and release in the cytosol, allowing the formation of an active BRI1-BAK1 receptor complex (18, 19). Additional mechanisms were coopted to stop BRI1 from firing after perception of BRs and allow plant cells to go back to the resting state. Autophosphorylation of residue S891 in the G-loop occurs late after BR perception and deactivates BRI1 via inhibiting its ATP binding (20). BRI1 is also subjected to internalization from the cell surface and vacuolar degradation using several mechanisms. First, the KINASE-ASSOCIATED PROTEIN PHOSPHATASE (KAPP) was proposed to regulate BRI1 through interaction between its forkhead-associated (FHA) domain and BRI1’s cytoplasmic domain (21). KAPP also co-localizes with the BAK1-related coreceptor SOMATIC EMBRYOGENESIS RECEPTOR KINASE1 (SERK1) at the plasma membrane and interacts with SERK1 in endosomes suggesting that KAPP-mediated dephosphorylation of BRI1 and SERK1 downregulates BR signaling (22). Second, BRI1 degradation was shown to require PROTEIN PHOSPHATASE2A (PP2A)-mediated dephosphorylation triggered by methylation of the PP2A using a leucine carboxylmethyltransferase (23). Most importantly, BRI1 undergoes endocytosis and degradation in the vacuole (24). This is controlled by lysine(K)-63 linked polyubiquitin chain conjugation to BRI1 intracellular domain driven by the PLANT U-BOX12 (PUB12) and PUB13 E3 ligases (25, 26). BRI1 ubiquitination promotes BRI1 internalization from the cell surface and is essential for proper sorting in endosomes and vacuolar targeting (25). BRI1 endocytosis was initially thought to be independent of ligand binding (24). However, the fact that BRI1 ubiquitination is dependent on both BRI1 kinase activity and ligand perception suggests that BRI1 internalization and vacuolar degradation is regulated by BRs (25, 26). BRI1 endocytosis is also under the control of environmental signals that impinge on growth via BR responses. Elevation of ambient growth temperature decrease BRI1 protein accumulation to boost heat-driven root elongation (27). The crosstalk between BR signaling and temperature responses likely uses BRI1 ubiquitination as the expression of an ubiquitination-defective BRI1 variant lacking 25 lysine residues in BRI1 intracellular domain abolishes BRI1 degradation upon warmth (27).

BRI1 serves as the archetypal plant receptor protein in the study of PTMs and their interplay, and their role in signaling/signal integration. Here show that BRI1 is SUMOylated and that BRI1-SUMO conjugates are regulated by the Desi3a SUMO protease. Most importantly, we uncover that BRI1 SUMOylation drops following growth at heightened temperatures, leading to increased BRI1 interaction with the negative regulator BIK1 and increased BRI1 endocytosis to attenuate BR-dependent growth. Finally, we demonstrate that such downregulation in BR signaling is required to restrict heat-induced growth responses. Overall, our work shed light on a new PTM targeting BRI1 and highlights its interplay with BRI1 ubiquitin-mediated endocytosis in the control of environmentally-regulated plant growth.

## RESULTS

### BRI1 is decorated with SUMO modifications *in planta*

BRI1 was previously demonstrated to be modified by ubiquitination on intracellular lysine residues (25). Ubiquitin is the founding member of a class of PTMs named Ubiquitin-Like modifiers (UBLs) that share a core β-grasp fold of approximately 70 amino acids and that can be reversibly attached to proteins or other cellular constituents lipids to regulate their activity (28). To evaluate if BRI1 is modified by other ubiquitin-like modifications, we first sought to monitor if the BRI1 is SUMOylated *in planta*. Transgenic plants expressing a functional BRI1 fusion to the mCitrine yellow fluorescent protein (mCit) under the control of *BRI1* promoter were used to immunoprecipitate BRI1-mCit protein with micromagnetic beads coupled to anti-GFP antibodies. Probing BRI1-mCit immunoprecipitates with anti-SUMO1 (AtSUMO1) revealed a SUMO specific signal at the size of BRI1-mCit, similar to what is observed for the positive controls JAZ6-GFP (29) (Figure 1A) or FLS2-GFP (Figure S1A) (30). This indicates that BRI1 is conjugated with SUMO1 under standard growth conditions. In contrast to the smear-like signals obtained for BRI1 with anti-Ub antibodies (25, 26), the SUMO1-specific signal associated to BRI1 migrated as a sharp band close to the molecular weight of BRI1 indicating that BRI1 carries a very limited number of SUMO modifications.

**Figure 1.**
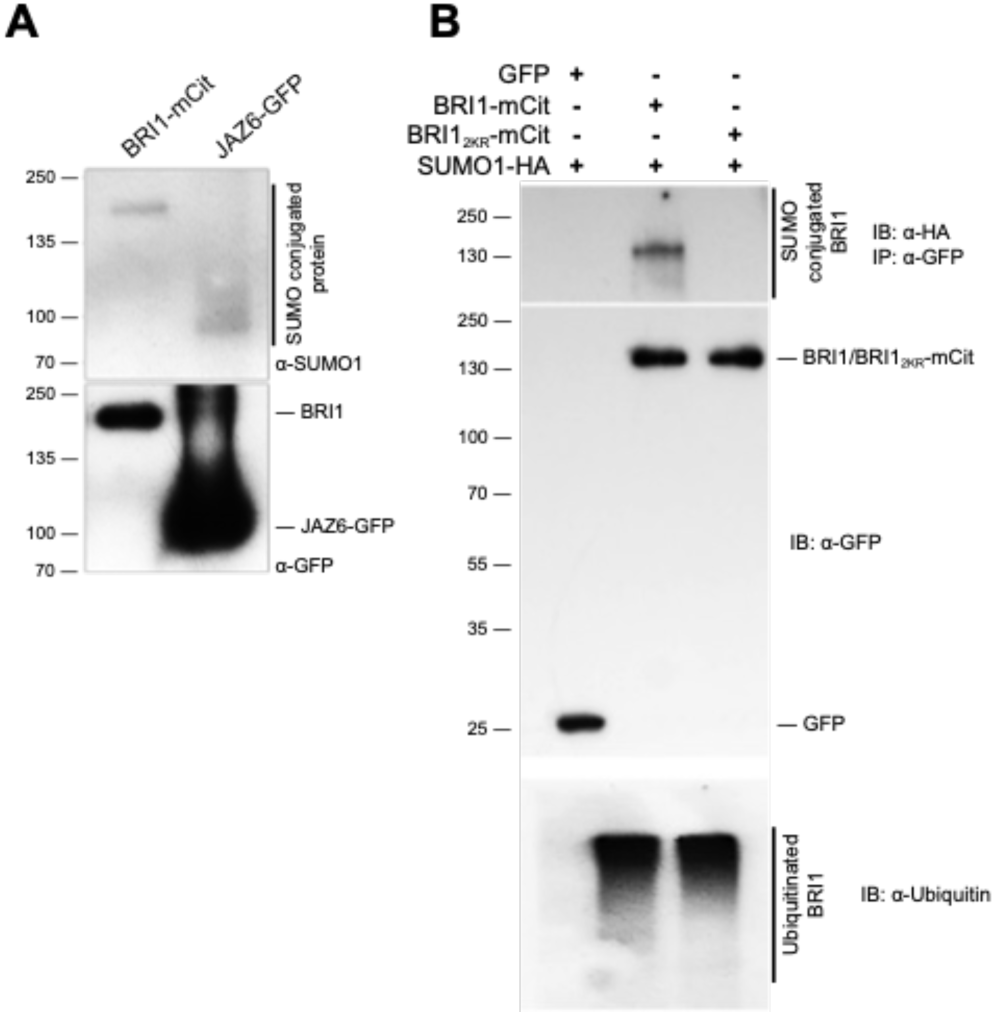
BRI1 is SUMOylated in vivo on two intracellular lysine residues. A. In vivo SUMOylation analyses of BRI1. Immunoprecipitation was carried out using anti-GFP antibodies on solubilized protein extracts from mono-insertional homozygous BRI1-mCitrine plants or the JAZ6-GFP positive control plants. Detection of immunoprecipitated proteins used the anti-GFP (bottom), and anti-SUMO1 (top) antibodies. B. In vivo SUMOylation analyses of BRI1 and BRI1_2KR_. BRI1-mCit and the BRI1_2KR_-mCit variant mutated for residues K1066,1118R were transiently expressed in *N. benthamiana* leaves prior to immunoprecipitation using anti-GFP antibodies on solubilized protein extracts. GFP alone was used as negative control. All constructs were co-expressed with SUMO1-HA. Detection of immunoprecipitated proteins used the anti-GFP (middle), anti-SUMO1 (top), and anti-ubiquitin (bottom) antibodies.

To gain further insight into BRI1 SUMOylation, we searched for possible SUMO sites in BRI1 as previously done with FLS2 (30). We identified lysine residues K1066 and K1118 in the BRI1 kinase domain as putative SUMO sites. Residue K1066 is strictly conserved in Arabidopsis BRI1-like proteins and more generally in plant BRI1 homologs (Figure S1B). A conserved lysine is also found in a close context to K1118 in Arabidopsis BRLs and plant BRI1 homologs. To decipher if both lysine residues are actual targets of SUMO *in planta*, we generated the K1066R and K1118R BRI1 variants where the corresponding lysine have been substituted to arginine to maintain the positive charge while preventing SUMOylation. Mutation of any of the two lysine residues in BRI1 decreased SUMO conjugation compared to wild-type BRI1 when transiently expressed in *Nicotiana benthamiana* (Figure S1C), suggesting that these are *bona fide* SUMO sites. Mutation of both lysine residues completely abolished BRI1 SUMOylation (Figure 1B), indicating that these are the only SUMO sites in BRI1. Mutation of K1066 and K1118 however did not alter significantly the overall ubiquitination pattern of BRI1 (Figure 1B). This observation is consistent with the fact that BRI1 is heavily ubiquitinated *in planta* and that mutation of 25 lysine residues in BRI1 intracellular domain, including K1066 and K1118, is required to completely abolish BRI1 ubiquitination (25). Altogether, our work reveals that BRI1 can be post-translationally modified at residues K1066 and K1118 by either ubiquitin or SUMO.

### BRI1 SUMOylation is regulated by the SUMO protease Desi3a

The levels of SUMO-conjugates of another plant plasma membrane protein, FLS2, are controlled by the balance between the SUMO E2-conjugating enzyme SCE1, which is capable of directly transferring SUMO onto target residues (31), and the Desi3a SUMO protease that deSUMOylates FLS2 (30). Desi3a has been shown to be degraded upon flagellin perception, allowing the accumulation of SUMO-FLS2 conjugates and triggering intracellular immune signaling. To examine if Desi3a also controls the levels of SUMO-BRI1, we first determined whether Desi3a is found in overlapping expression territories with BRI1. Publicly available genome-wide expression data reveals that *Desi3a* has a broad expression profile overlapping with *BRI1*. We confirmed these observations by RT-PCR using RNA extracted from various Arabidopsis tissues (Figure S2). We next addressed whether Desi3a protein co-localizes with BRI1 at the plasma membrane. Transient expression of a Desi3a-mCherry (Desi3a-mCh) fusion protein in *N. benthamiana* confirmed the presence of Desi3a at the cell surface (Figure 2A). A similar pattern was observed with the BRI1-mCit of FLS2-GFP functional fusions. In addition, a clear colocalization at the cell surface is observed between Desi3a and BRI1 or FLS2 (Manders coefficient M_BRI1_= 0.83 and M_FLS2_=0.80), to the resolution of the confocal microscope. The fluorescence profile of Desi3a-mCh clearly overlapped with BRI1-mCit at the plasma membrane, similarly to what is observed for FLS2-GFP (Figure S3) (30). We next tested whether the Desi3a SUMO protease had the ability to interact with BRI1. To this purpose, we took advantage of the transient expression in *N. benthamiana* since BRI1 is SUMOylated in this experimental system. BRI1-mCit was transiently expressed together with Desi3a-HA and subjected to immunoprecipitation using GFP beads. Probing BRI1-mCit immunoprecipitates with anti-HA antibodies revealed the presence of Desi3a-HA (Figure 2B). Expression of GFP alone with Desi3a was used as a negative control and failed to capture any interaction (Figure 2B), indicating that BRI1 interacts *in vivo* with Desi3a. Furthermore, we used the ULP1a SUMO protease that deSUMOylates the BZR1 transcription factor (32) and observed no interaction with BRI1 (Figure 2B).

**Figure 2.**
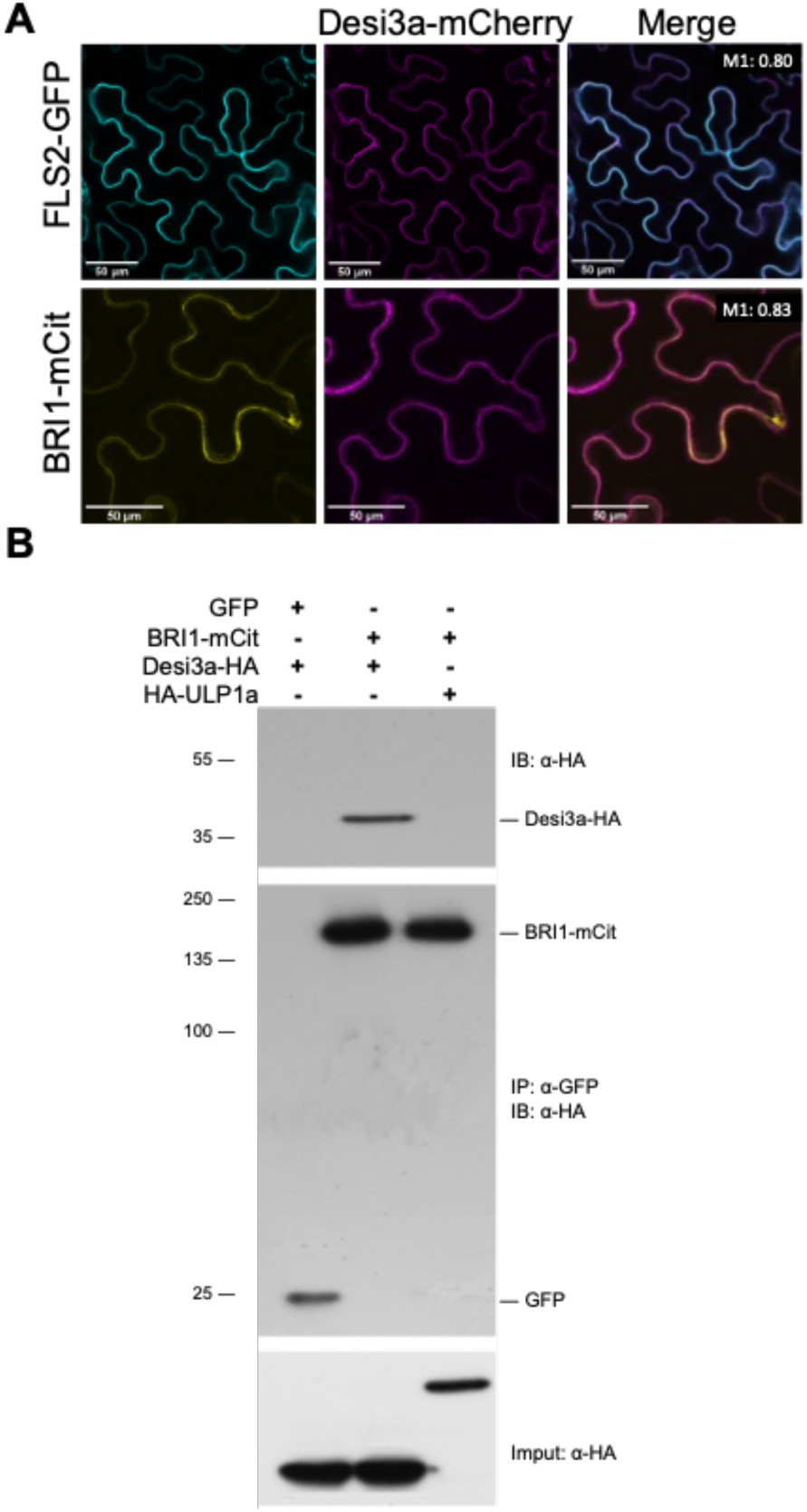
BRI1 interacts with the Desi3a SUMO protease in vivo. A. Confocal microscopy analyses and colocalization analyses between BRI1 and Desi3a. BRI1-mCit and Desi3a-mcherry (Desi3a-mCh) were transiently expressed in *N. benthamiana* leaves. The Manders colocalization coefficient are shown in the overlay channel. Scale bars=50 μm. B. In vivo interaction between BRI1 and Desi3a. BRI1-mCit and Desi3a-HA were transiently expressed in *N. benthamiana* leaves prior to immunoprecipitation using anti-GFP antibodies on solubilized protein extracts. GFP alone and the ULP1a SUMO protease were used as negative control. Detection of immunoprecipitated proteins used the anti-GFP (middle) and anti-HA (top) antibodies. The input faction of Desi3a-HA and HA-ULP1a is shown at the bottom.

Altogether, this indicates that BRI1 specifically interacts with the plasma membrane-localized Desi3a SUMO protease and that the SUMOylation status of BRI1 is controlled by the Desi3a SUMO protease.

### Temperature elevation regulates BRI1 deSUMOylation

To shed light on the biological relevance of BRI1 SUMOylation, we searched for possible stimuli/conditions increasing or decreasing the accumulation of BRI1 SUMO conjugates. Among conditions tested, we focused our attention on the role of ambient temperature elevation since previously reported to post-translationally regulate BRI1 (27). Plants expressing BRI1-mCit were grown at 21°C or 26°C, as previously done (27), before immunoprecipitation of BRI1-mCit. Immunoprecipitates were normalized to show equivalent BRI1-mCit signals, as visualized using anti-GFP antibodies (Figure 3A), before being probed with anti-SUMO1 antibodies to detect the SUMOylated pool of BRI1 at both temperatures. Plants grown at 21°C or 26°C clearly showed different accumulation of BRI1 SUMO conjugates. Notably, temperature elevation was accompanied with lower SUMOylated BRI1 (Figure 3A). Quantification of the BRI1-SUMO signals obtained, relative to the levels of immunoprecipitated BRI1 revealed a 7-fold decrease at elevated temperature (Figure 3B). Such a drop in BRI1-SUMO levels at higher temperature may be a direct consequence of increased Desi3a levels. To test this, we first investigated the influence of temperature on the accumulation of BRI1 and Desi3a proteins using transient expression in *N. benthamiana*. Plants agroinfiltrated with BRI1-mCit and Desi3a-mCh were exposed to either 21°C or 26°C for 2 days and imaged at the confocal microscope. Several independent experiments were carried out and multiple regions of interest analyzed to overcome the variability in transformation efficiency. Overall, temperature elevation reproducibly decreased BRI1-mCit accumulation when co-expressed with Desi3a-mCh (Figure 3C, 3D), reminiscent of the effect of heat on BRI1 accumulation previously observed in Arabidopsis roots (27). This was also accompanied by an increase in the fluorescence associated with Desi3a (Figure 3C, 3D). The impact of temperature elevation was confirmed using transgenic Arabidopsis plants constitutively expressing a Desi3a-HA fusion protein. Heat promoted the accumulation of Desi3a-HA protein as observed by western blot (Figure 3E, F), pointing to a post-transcriptional regulation of Desi3a by temperature. We then addressed how temperature affects the ability of BRI1 and Desi3a to interact *in vivo*. Agroinfiltrated plants exposed to 26°C reproducibly harbored lower BRI1-mCit accumulation (Figure 3G), consistent with our confocal microscopy observations. BRI1 immunoprecipitates however showed increased Desi3a-HA proteins levels at 26°C, pointing to the increased interaction between BRI1 and Desi3a when plant experience heightened temperature (Figure 3G). Taken together, these observations indicate that the increased accumulation of Desi3a and stronger interaction between BRI1 likely explains the drop in BRI1 SUMOylation observed at 26°C.

**Figure 3.**
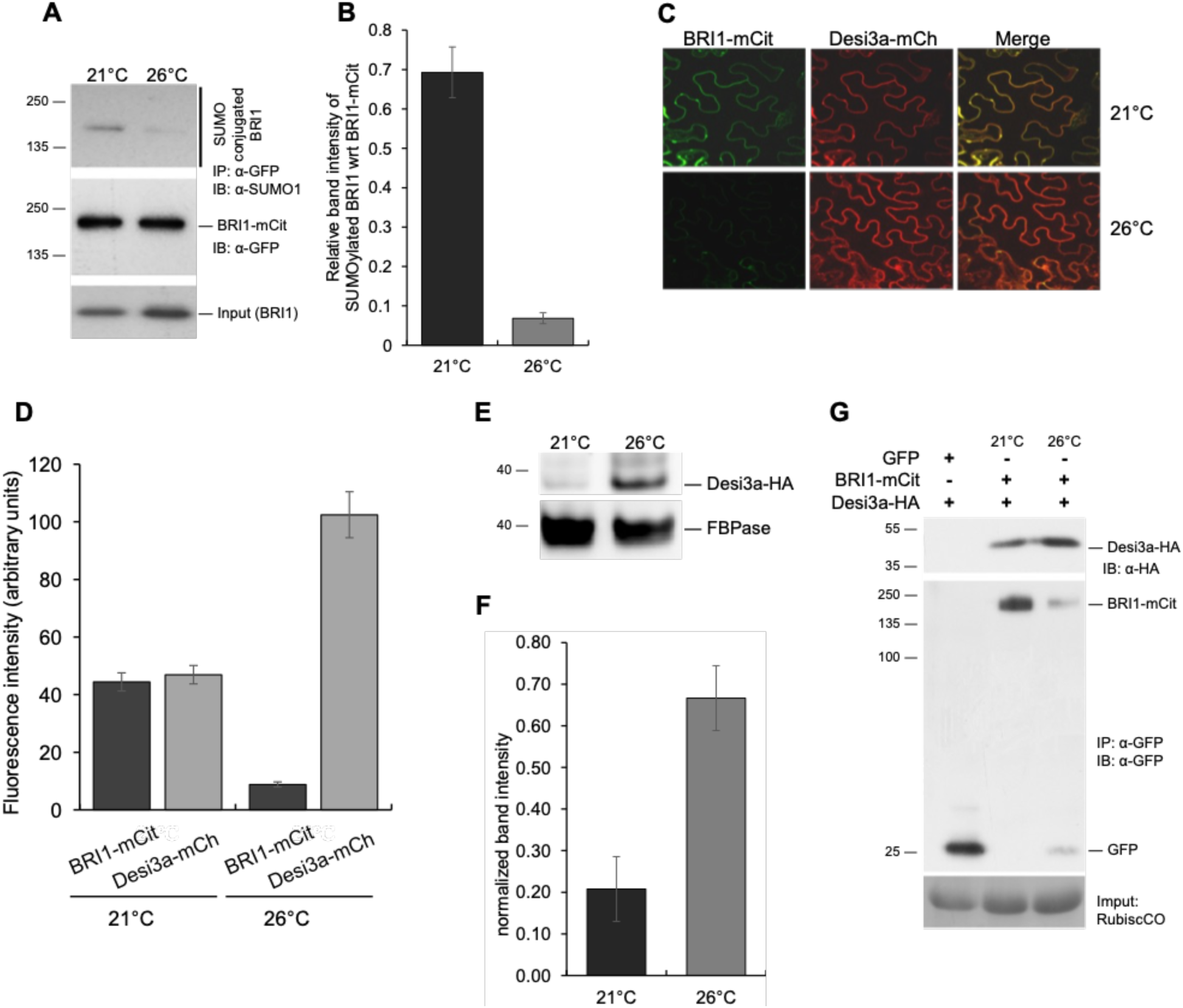
Desi3a deSUMOylates BRI1 upon changes in ambient temperature. A. In vivo SUMOylation analyses of BRI1 at 21°C or 26°C. Immunoprecipitation was carried out using anti-GFP antibodies on solubilized protein extracts from mono-insertional homozygous BRI1-mCitrine plants grown at 21°C or 26°C. Detection of immunoprecipitated proteins used the anti-GFP (middle) and anti-SUMO1 (top) antibodies. The input fraction for BRI1-mCit is shown at the bottom. B. Quantification of BRI1 SUMO at 21°C and 26°C relative to immunoprecipitated BRI1-mCit levels. Error bars represent SEM (n=3). C. Confocal microscopy analyses of BRI1 and Desi3a at 21°C and 26°C. BRI1-mCit and Desi3a-mch were transiently expressed in *N. benthamiana* leaves and incubated at 21°C or 26°C for 2 days. Similar confocal detection settings were used to compare the effect of temperature on BRI1 and Desi3a proteins levels. Several independent experiments were carried out and multiple regions of interest analyzed to overcome the variability in transformation efficiency. Scale bars=20 μm. D. Quantification of BRI1 and Desi3a fluorescence levels in experiments carried out as in C. Multiple regions of interest analyzed for to overcome the variability in transformation efficiency. Error bars represent SEM (n=15). The asterisk indicates a statistically significant difference in BRI1-SUMO at 26°C (Mann-Whitney). E. In vivo interaction between BRI1 and Desi3a at 21°C and 26°C. BRI1-mCit and Desi3a-HA were transiently expressed in *N. benthamiana* and incubated at 21°C or 26°C for 2 days prior to immunoprecipitation using anti-GFP antibodies on solubilized protein extracts. GFP alone was used as negative control. Detection of immunoprecipitated proteins used the anti-GFP (middle) and anti-HA (top) antibodies. Ponceau staining showing RubisCo accumulation is used as loading control. F. Western blot analyses monitoring the accumulation of Desi3a-HA protein in plants grown at 21°C or 26 °C. Detection of Desi3a-HA is performed with anti-HA antibodies. The membrane was stripped and probed with anti-FBPase antibodies as loading control. G. Quantification of Desi3a protein at 21°C and 26°C relative to FBPase levels. Error bars represent SEM (n=3). The asterisk indicates a statistically significant difference in BRI1-SUMO at 26°C (Mann-Whitney).

### Desi3a-mediated deSUMOylation of BRI1 controls temperature responses

Plant responses to temperature are highly dependent on other environmental factors such as light (27, 33). We therefore assessed the genetic contribution of BR signaling to plant responses to elevated temperature in our conditions by scoring hypocotyl length of wild-type, *bri1*, and the *bes1-D* constitutive BR response mutant at 21°C or 26°C. Wild-type seedlings grown at elevated temperature elongated their hypocotyls (Figure S4A, S4B, S4C). *bes1-D* showed much greater responses to heat than wild-type while *bri1* failed to respond (Figure S4A, S4B, S4C), suggesting that BR signaling positively impinges on plant temperature responses in hypocotyls. We next investigated the role of SUMO/deSUMOylation upon warmth by comparing hypocotyl length of wild-type, and *desi3a* at 21°C or 26°C. *desi3a* mutants showed slightly shorter hypocotyls at 21°C compared to wild-type seedlings, but elongated more at 26°C (Figure 4A). This increased response to temperature is clearly highlighted by the elevated ratio of hypocotyl length at 26°C/21°C for *desi3a* (Figure 4B). This suggests that Desi3a is a negative regulator of temperature responses or that conversely, SUMOylation is required to promote hypocotyl elongation upon heat. We next monitored the phosphorylation status of the BR pathway downstream transcription factor BR BES1. BES1 exists under a phosphorylated form (P-BES1) and dephosphorylated form (BES1), and exogenous BR application promotes the conversion of P-BES1 into its unphosphorylated active BES1 form (11). Compared to wild-type plants, *desi3a* showed a mild increase in dephosphorylated BES1:phosphorylated BES1 ratio levels (Figure 4C, 4D), indicating that *desi3a* harbors enhanced BR signaling. This observation is consistent with *bes1-D* greatly over-responding to high temperature (Figure S4A, S4B, S4C).

**Figure 4.**
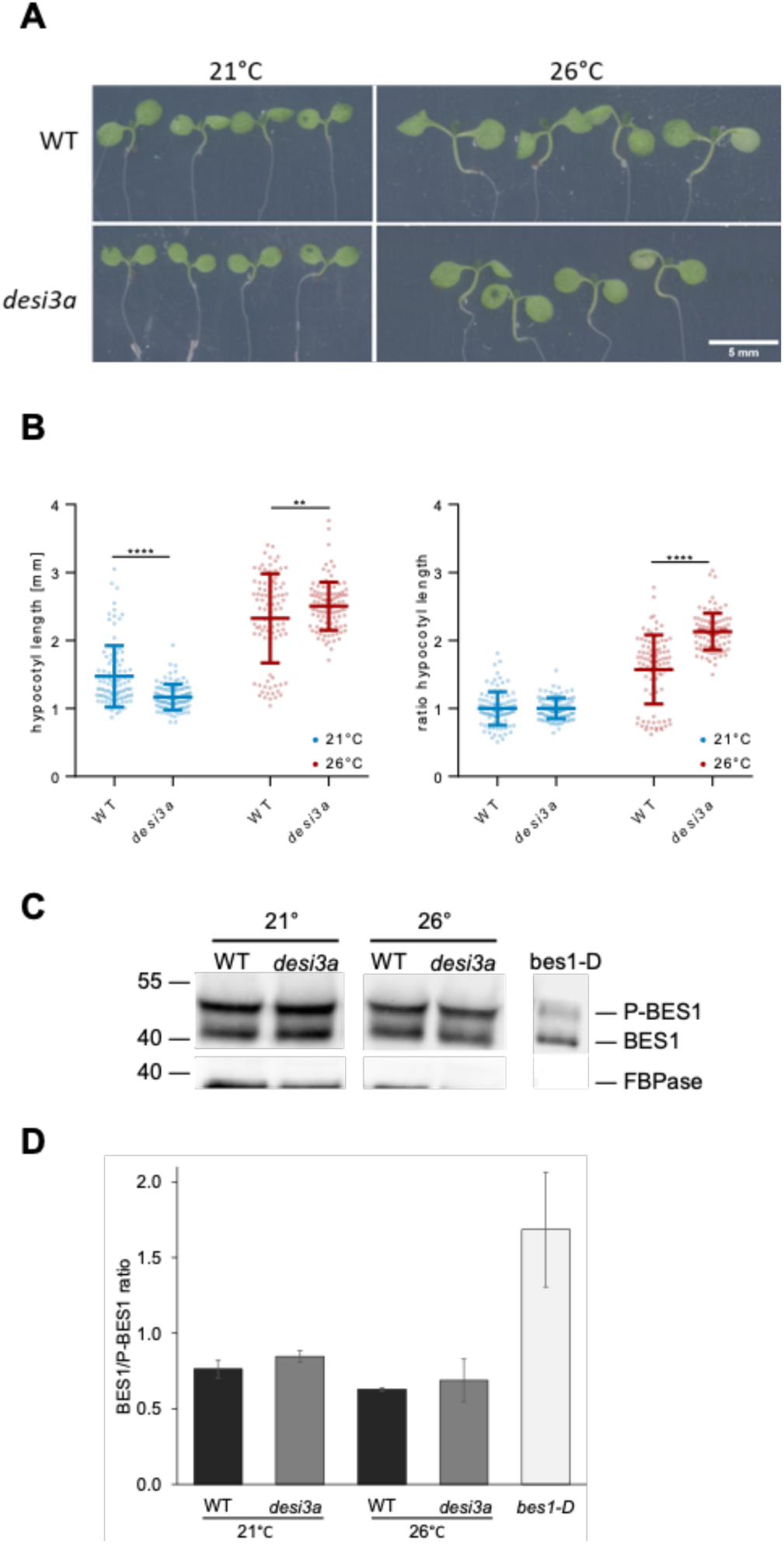
Desi3a is a negative regulator of plant responses to temperature elevation. A. Phenotype of 6-day-old wild-type (WT) and *desi3a* mutant plants grown at 21°C or 26°C. Representative pictures are shown. B. Hypocotyl length of 6-day-old wild-type (WT) and *desi3a* mutant plants grown at 21°C or 26 °C. Experiments were carried out in triplicates. Error bars represent SEM (n=20). The asterisk indicates a statistically significant difference between WT and *desi3a* (two-way ANOVA with Sidak’s multiple comparison test). C. Ratio of hypocotyl length from wild-type (WT) and *desi3a* mutant plants grown at 21°C and 26 °C for 6 days. Experiments were carried out in triplicates. Error bars represent SEM (n=20). The asterisk indicates a statistically significant difference between wild-type and *desi3a* (two-way ANOVA with Sidak’s multiple comparison test). D. Phosphorylation state of the BES1 transcription factor in wild-type (WT) or *desi3a* plants grown at 21°C or 26 °C. Detection of BES1 is performed with anti-BES1 antibodies. E. Quantification of the ratio between BES1 and phosphorylated BES1 (P-BES1). Error bars represent SEM (n=2). The asterisk indicates a statistically significant difference in BRI1-SUMO at 26°C (Mann-Whitney).

Next, we sought to decipher the specific role of BRI1 SUMO/deSUMOylation in plant responses to elevated temperature and the underlying mechanism(s) by genetically impacting on BRI1 SUMOylation. We reasoned that generating transgenic plants expressing the non-SUMOylated BRI1_2KR_ version would prevent us from reaching any solid conclusion on the role of BRI1 SUMO/deSUMOylation since K1066 and K1118 are also ubiquitin targets. Mutating both lysine residues would indeed directly abolish ubiquitination and SUMOylation at these sites, and possibly other lysine-based post-translational modifications. We decided instead to characterize deeper the impact of altered BRI1-SUMO levels at residues K1066 and K1118 using the *desi3a* mutant background. This offers the great advantage to grasp the interplay between both post-translational modifications at K1066 and K1118. We therefore crossed the *desi3a* T-DNA knock-out mutant with the BRI1-mCit reporter line and isolated *desi3a*/BRI1-mCit double homozygous plants. These plants were imaged at the confocal microscope to observe any possible change in BRI1 distribution in the cell. No obvious change in BRI1 distribution between the plasma membrane in endosomes of BRI1-mCit or *desi3a*/BRI1-mCit lines. To better investigate the possible effect of BRI1 SUMO on BRI1 dynamics, we then took advantage of the fungal toxin Brefeldin A (BFA) that inhibits endosomal trafficking in Arabidopsis roots and hypocotyls (34, 35), and thus creates large aggregates of *trans*-Golgi network/early endosomal compartments. Endocytosed BRI1-mCit was found in BFA bodies when plants were challenged with BFA in the presence of the translation inhibitor cycloheximide (CHX) (Figure 5A), similar to roots (24, 25, 36). However, quantification of several parameters pointed to a reduction in BRI1 endocytosis in the *desi3a* mutant compared to wild-type plants. First, the ratio of plasma membrane over intracellular BFA-trapped BRI1-mCit fluorescence is increased in *desi3a* compared to the wild-type background (Figure 5B). Second, loss of Desi3a is accompanied with a reduction in the number of BFA bodies per cell (Figure 5C). Taken together, these observations indicate the loss of Desi3a and SUMOylation at residues K1066 and K1118 directly or indirectly decreased the endocytic flux of BRI1.

**Figure 5.**
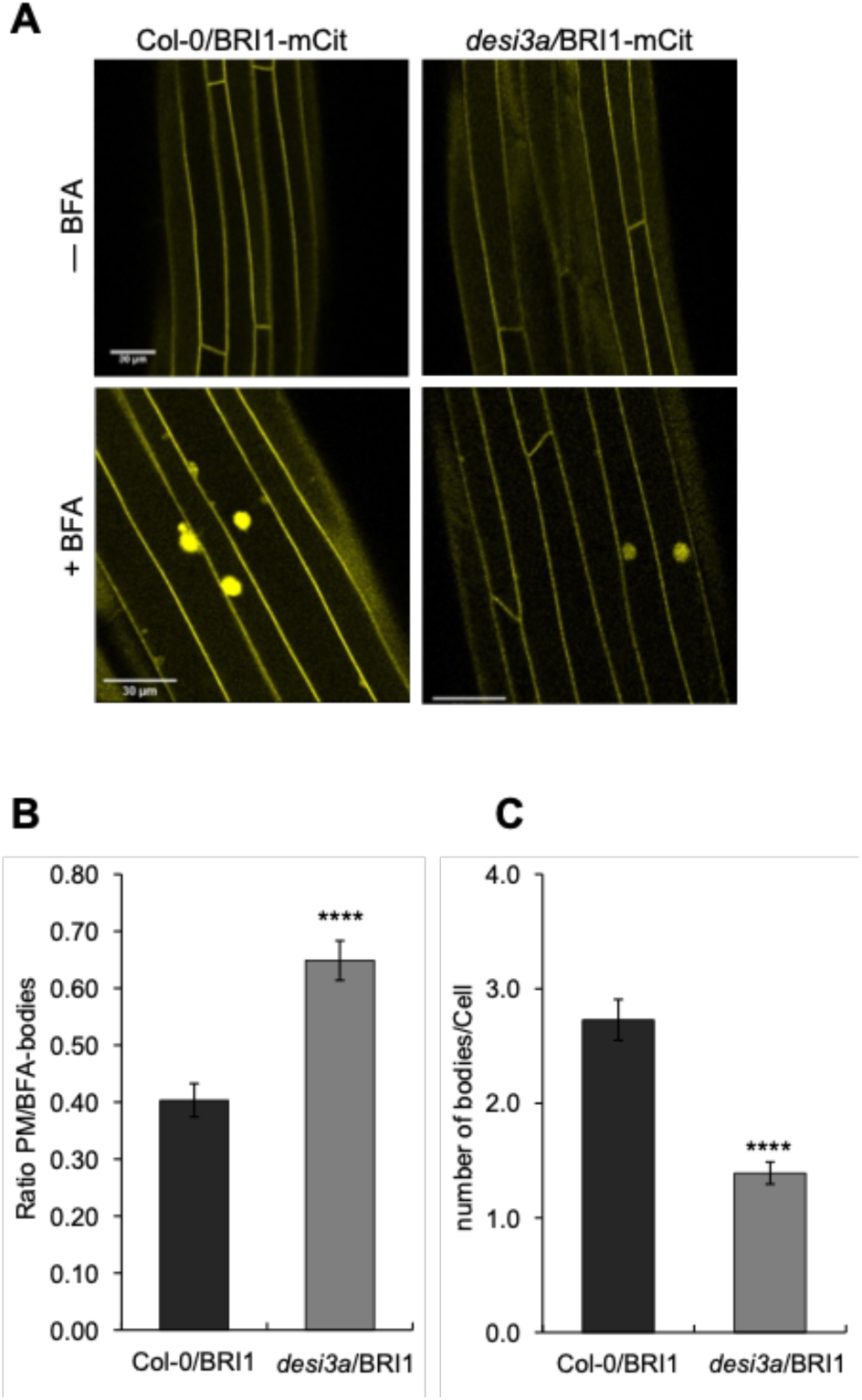
Desi3a-dependent deSUMOylation regulates BRI1 endocytosis and interaction with BIK1. A. Confocal microscopy analyses of BRI1-mCit and *desi3a*/BRI1-mCit dark-grown hypocotyls. Similar confocal detection settings were used to compare the fluorescence intensity in the two transgenic lines. Scale bars=30 μm. B. Quantification of the ratio between plasma membrane and BFA body fluorescence signal intensities of BRI1-mCit and *desi3a*/BRI1-mCit. C. Quantification of the number of BFA bodies per cell in BRI1-mCit and *desi3a*/BRI1-mCit. Experiments were carried out in triplicates. Error bars represent SEM (n=9). The asterisk indicates a statistically significant difference between BRI1-mCit and *desi3a*/BRI1-mCit (Mann-Whitney).

### BIK1 is recruited to deSUMOylated BRI1 to dampen temperature responses

Lack of SUMOylation renders FLS2 unable to mount proper immune responses due to the increased interaction with the downstream receptor-like cytoplasmic kinase (RLCK) BIK1 (30), which acts as a positive regulator of FLS2-mediated signaling (37). BIK1 is also known to participate to BR signaling, although as negative regulator, where it is associated with BRI1 in resting cells and is released from the BRI1-BAK1 receptor complex upon BR perception (38). To shed light on the direct functional consequences of BRI1 SUMO/deSUMOylation, we therefore investigated if BRI1 SUMOylation affects the BRI1-BIK1 interaction using transient expression in *N. benthamiana*. Immunoprecipitation of BIK1-HA using HA beads followed by probing with anti-GFP antibodies failed to detect any interaction with free GFP (Figure 6A). In contrast, BIK1-HA was able to interact with BRI1-mCit (Figure 6A), consistent with previous reports (38). Strikingly, BIK1-HA showed a much stronger interaction with the SUMO-defective BRI1_2KR_ (Figure 6A). Considering that BRI1 SUMO levels are lower at 26°C, we sought to determine the influence of temperature elevation on the BRI1-BIK1 interaction. BRI1 immunoprecipitates recovered higher BIK1 protein levels at 26°C compared to 21°C (Figure 6B), consistent with the fact that BIK1 shows stronger interaction with the non-SUMOylatable BRI1_2KR_ variant. This evidence indicates that, similarly to what was observed for FLS2 (30), SUMOylation reduces the interaction of BRI1 with the downstream kinase BIK1. Genetically, BIK1 negatively regulates BR signaling, with *bik1* mutant showing increased BR responses even upon high concentration of the BRZ BR biosynthetic inhibitor (38). To further illustrate the contribution of BRI1 SUMOylation to temperature responses and determine the possible role of BIK1 in this process, we phenotyped the previously published *bik1* loss-of-function mutant at 21°C and 26°C. In contrast to the mild increase in hypocotyl length observed in wild-type seedlings, *bik1* hypocotyls dramatically elongated in response to heat (Figure 6C, 6D, 6E). These observations point to the role of BIK1 as negative regulator of temperature responses, similarly to Desi3a.

**Figure 6.**
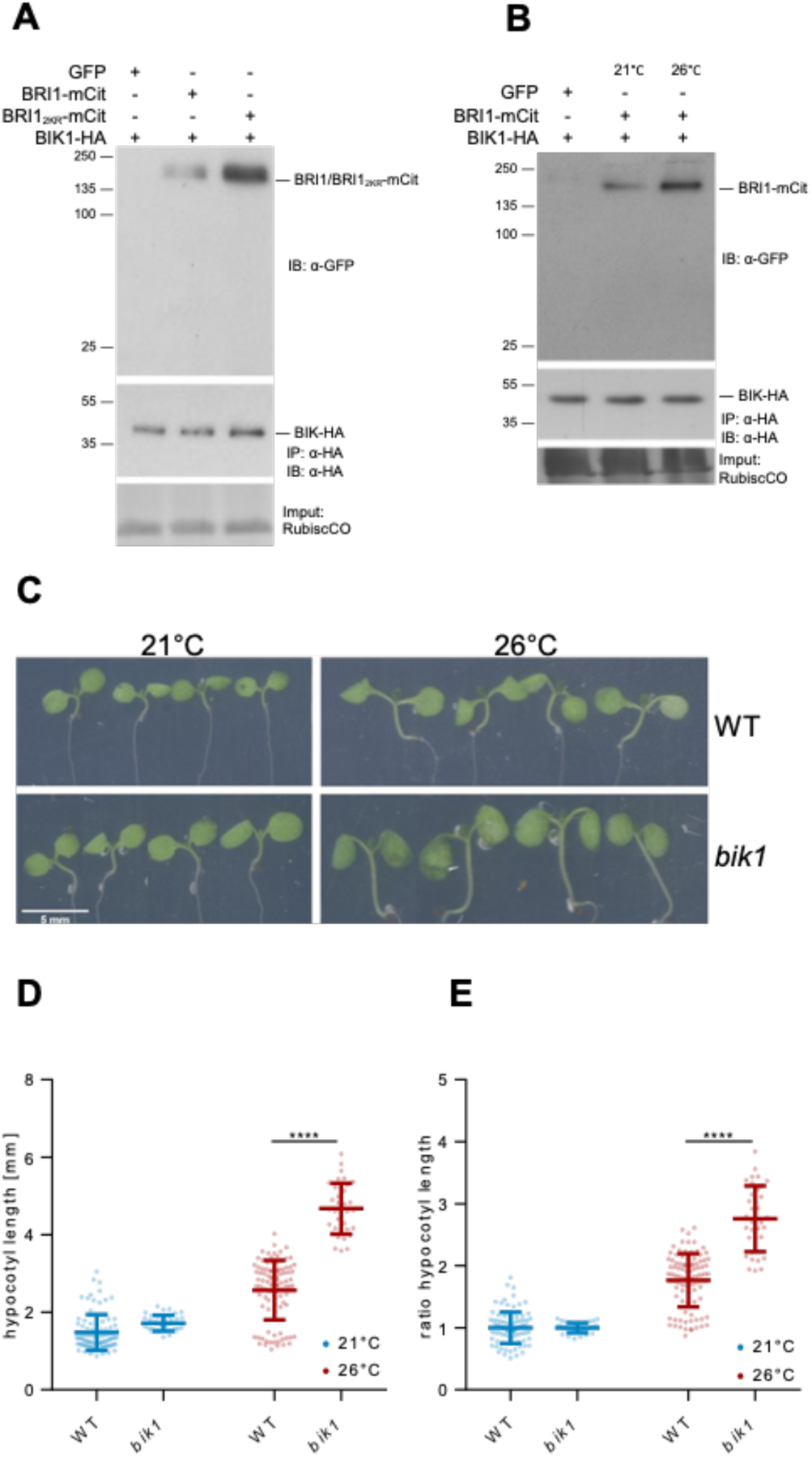
Desi3a-dependent deSUMOylation regulates BRI1 interaction with BIK1. A. In vivo interaction between BIK1 and wild-type BRI1 or BRI1_2KR_. BIK1-HA was coexpressed with BRI1-mCit or BRI1_2KR_-mCit in *N. benthamiana* leaves prior to immunoprecipitation using anti-HA antibodies on solubilized protein extracts. GFP alone was used as negative control. Detection of immunoprecipitated proteins used the anti-HA (middle) and anti-GFP (top) antibodies. Ponceau staining showing RubisCo accumulation is used as loading control. B. In vivo interaction between BIK1 and BRI1 at 21°C or 26°C. BIK1-HA and BRI1-mCit were transiently expressed in *N. benthamiana* and incubated at 21°C or 26°C for 2 days prior to immunoprecipitation using anti-HA antibodies. GFP alone was used as negative control. Detection of immunoprecipitated proteins used the anti-HA (middle) and anti-GFP (top) antibodies. Ponceau staining showing RubisCo accumulation is used as loading control. C. Phenotype of 6-day-old wild-type (WT) and *bik1* mutant plants grown at 21°C or 26°C. Representative pictures are shown. D. Hypocotyl length of 6-day-old wild-type (WT) and *bik1* mutant plants grown at 21°C or 26 °C. Experiments were carried out in triplicates. Error bars represent SEM (n=20). The asterisk indicates a statistically significant difference between WT and *bik1* (two-way ANOVA with Sidak’s multiple comparison test). E. Ratio of hypocotyl length from wild-type (WT) and *bik1* mutant plants grown at 21°C and 26 °C for 6 days. Experiments were carried out in triplicates. Error bars represent SEM (n=20). The asterisk indicates a statistically significant difference between wild-type and *bik1* (two-way ANOVA with Sidak’s multiple comparison test).

## DISCUSSION

The BR receptor BRI1 serves as a model to grasp the intricate mechanisms of RLK-mediated signaling in plants (39). The activity of BRI1 is regulated by post-translational modifications required to initiate, amplify or dampen BR signaling. Phosphorylation between BRI1, the SERK coreceptors, and different downstream receptor-like cytoplasmic kinases mostly activates BR signaling (39), while ubiquitination leads to BRI1 degradation and signal attenuation (25, 26). Several endogenous and exogenous cues also converge at the level of BRI1 protein to influence BR-dependent growth. For example, glucose was shown to increase BRI1 endocytosis to control, in part, changes in root architecture associated with fluctuations in light intensity and photosynthetic activity (40). Similarly, elevation of ambient temperature impinges on BR-dependent root growth by triggering BRI1 destabilization (27). We show here that BRI1 is subjected to SUMO/deSUMOylation and that this is also required to control plant responses to heat. BRI1 is decorated with SUMO under standard conditions, and deSUMOylated by the Desi3a SUMO protease when plants are grown at warm temperature (Figure S5). Desi3a-mediated BRI1 deSUMOylation favors BRI1 interaction with the negative regulator of BR signaling BIK1 and also promotes BRI1 endocytosis. *desi3a* and *bik1* mutants both overrespond to heightened temperature, indicating that both Desi3a and BIK1 act as negative regulators of temperature responses. Thus, our work highlights a new mechanism to integrate temperature input into BR-dependent growth responses.

The major driver of the elongation observed when plants face heat is the phytohormone auxin (41). Temperature elevation reduces PHYTOCHROME B activity and induces *PHYTOCHROME INTERACTING FACTOR4* (*PIF4*) expression to increase auxin biosynthesis (42–49). PIF4, as well as other PIFs, directly binds to the promoters of auxin biosynthesis genes, such as *YUCCA8* (*YUC8*) and *YUC9*, *TRYPTOPHAN AMINOTRANSFERASE OF ARABIDOPSIS1* (*TAA1*) and *CYTOCHROME P450 FAMILY79B* (*CYP79B2*) to increase auxin-responsive gene expression and tissue elongation (42, 43, 45, 47–50). BRs were also reported to participate to heat response in both aerial and underground tissues, although their contribution is opposite in both organs where BRs are known to differentially regulate growth (14, 51). BZR1 binds to the promoter of *PIF4* at elevated temperature to increase its expression and amplify shoot transcriptional responses to heat and promotes hypocotyl elongation (52). In roots, heat promotes BRI1 destabilization to downregulate BR signaling and to stimulate root elongation (27). Our findings now shed light on another level of control of BR signaling by temperature in aerial parts, with heat promoting BIK1 recruitment to the BR receptor complex and decreasing BRI1 levels through Desi3a-mediated BRI1 deSUMOylation (Fig. S5). This new layer of integration between BR signaling and warmth negatively regulates temperature responses, as loss of Desi3a or BIK1 yields increased heat-induced hypocotyl elongation. Auxin and BRs are well-known to act synergistically to promote cell elongation (53). The SUMO-dependent regulation of BRI1 we have uncovered likely allows plants to dampen BR signaling and to balance the synergistic effect between auxin and BRs on the transcription of growth-promoting genes, thus preventing plants from over-elongating upon warmth. The precise molecular mechanisms underlying this new regulatory level are still unclear. SUMOylation may have a direct inhibitory role on BRI1 endocytosis so that Desi3a-triggered BRI1 deSUMOylation at 26°C increases internalization of BRI1. There are few reports describing a role for SUMO in inhibiting endocytosis. For example, SUMOylation of the TRPM4 Ca^2+^-activated nonselective cation channel impairs TRPM4 endocytosis and leads to elevated TRANSIENT RECEPTOR POTENTIAL CATION CHANNEL SUBFAMILLY M4 (TRPM4) density at the cell surface (54). Alternatively, SUMO/deSUMOylation of BRI1 may indirectly impact on BRI1 dynamics *via* the interplay between SUMO and Ub in the control of BRI1 endocytosis. BRI1 internalization from the cell surface is driven by massive ubiquitination decorating many cytosol-exposed lysine residues through the PUB12 and PUB13 E3 Ub ligases (25, 26). Additionally, heat was shown in roots to destabilize BRI1 in an ubiquitin-dependent manner (27). Similarly to Ub, SUMO is covalently attached to proteins using lysine indicating that SUMO modifications potentially compete with Ub for the same sites to alter protein functions (55). During NF-κB activation for example, IκBα is ubiquitinated and degraded to release its inhibition of NF-κB (56).

In contrast, SUMOylation at the same site prevents ubiquitination and turnover of IκBα (57), therefore inhibiting NF-κB activation. BRI1 SUMOylation at residues K1066 and K1118 may therefore limit its ubiquitination at standard growth temperature. The deSUMOylation of BRI1 observed at elevated temperature and driven by Desi3a would free additional lysine residues for ubiquitination, thus boosting Ub-mediated endocytosis of BRI1. No significant difference in the Ub profile of BRI1 could however be observed in the SUMO-defective BRI1_2KR_ form. The presence of many target lysines for Ub in BRI1 yields a large high molecular weight BRI1-Ub smear that likely masks the effect of lack of ubiquitination at K1066 and K1118 in BRI1_2KR_. Consistently, the mutation of a single Ub target lysine in BRI1 identified by proteomics had no significant impact of the overall Ub profile of BRI1 (25). Considering the prominent role of Ub in endocytosis of plasma membrane proteins, we propose that heat-regulated SUMO/deSUMOylation of BRI1 allows plant to fine tune BRI1 Ub-mediated endocytosis and BR signaling (Fig. S5). Whether PUB12 and PUB13 are responsible for the linkage of additional Ub chains to deSUMOylated BRI1 upon heat or whether a yet to be characterized temperature-regulated E3 Ub ligase is involved will have to be tackled in the future.

The FLS2 LRR-RLK flagellin receptor shows increased SUMOylation upon flagellin perception mediated by Desi3a degradation, thus releasing BIK1 from FLS2 (30). BIK1 being a positive regulator of FLS2 signaling, FLS2-SUMO conjugates positively regulate downstream innate immune signaling (30, 37). BRI1 and FLS2 are often compared since both representing model for the large plant LRR-RLK family. Despite their radically different biological outputs, BRI1- and FLS2-mediated signaling pathways share striking parallels. Both receptors predominantly localize at the plasma membrane where they bind their respective ligands and fire (58, 59). BRI1 and FLS2 both use the same subset of co-receptors to initiate signaling (8, 9, 60, 61). Structural and biochemical studies revealed that BRs and the flg22 flagellin peptide both act as molecular glue to form or stabilize signaling-competent receptor complexes (3, 6, 62). In both cases, ligand binding triggers *cis*- and *trans*-phosphorylation events within the receptor complexes to reach full activation (63, 64). Downstream of the receptor complexes are found RLCKs that are direct substrates of the receptor complexes (65). In particular, the RLCK BIK1 is shared between both pathways but is a positive regulator for immune responses and a negative regulator for BR signaling (37, 38, 66). BIK1 has been shown to negatively regulate BR signaling through direct association with BRI1 (38). After BR perception, BIK1 is phosphorylated by BRI1, causing its dissociation from the receptor. The precise function of BIK1 in BR signaling is therefore still unclear. Regardless, the Desi3a- and SUMO-regulated interaction of BIK1 with BRI1 and FLS2 now emerges as a possible common regulatory mechanisms of ligand-binding LRR-RLs and corresponding signaling pathways. Although BRI1 and FLS2 have been shown to be confined to different nanodomains at the plasma membrane and the corresponding pathways to use different phosphocode in early phases (67), how signaling specificity is maintained along both pathways that rely on several shared components will deserve more attention in the future.

## MATERIAL & METHODS

### Plant material and growth conditions

The genotypes used in this study are wild-type (Col0), *bri1* T-DNA knockout (GABI_134E10) (19), Desi3a-HA (30), *desi3a-1* T-DNA knockout (SALK_151016C) (30), *bik1* T-DNA knock-out (SALK_005291) (37), *bes1-D* (11), FLS2::FLS2-GFP (30) and 35S::JAZ6-GFP (29). BRI1_K1066R_, BRI1_K1118R_ and the BRI1_2KR_ variant carrying the K1066R and K1118R substitutions were generated by site-directed mutagenesis of the pDONR221-BRI1 (19) using the primers listed (Table S1). Final destination vectors obtained by recombination using the pB7m34GW destination vectors (68), and the entry vectors pDONRP4P1r-BRI1prom (19), pDONR221-BRI1 or mutated BRI1 versions, and pDONRP2rP3-mCitrine (19).

After seed sterilisation and stratification, seeds were placed for germination on solid agar plates containing half-strength Linsmaier and Skoog medium without sucrose. They were grown vertically in growth chambers under long-day conditions (16 h light/8 h dark, 90 µE m^-2^×s^-1^) at 21°C or 26°C. For the specific growth conditions, refer to figure legends.

Infiltration of *N. benthamiana* leaves was performed using standard procedures and used the binary vectors carrying BRI1::BRI1-mCit (19), 35S::Desi3a-mCh and FLS2::FLS2-GFP (30).

### Chemical treatments

The final concentrations of the chemicals are indicated in the figure legends. The chemical stock solutions are in the following concentration: 100 mM cycloheximide (Sigma) in EtOH, 10 mM BFA (Sigma) in DMSO.

### Hypocotyl measurements

Seeds were sterilised and stratified for four-days. For dark-grown hypocotyl measurements, seeds were exposed to the light for 6h and then placed in dark at 21°C or 26°C for 3 days. For light-grown hypocotyl measurements, seeds were directly exposed to light at 21°C or 26°C for 6 days. Plates were scanned and hypocotyls measured using Fiji imageJ software. The mean and standard error to the mean (SEM) were calculated by combining the three replicates.

### Immunoprecipitation and western Blot analysis

For detection of proteins from crude extracts, total proteins were extracted from ∼50 mg plant material using Laemmli extraction buffer, using a 1:3 w/v ratio between tissue powder and extraction buffer. After debris elimination, proteins were separated by SDS-PAGE. Protein detection was carried out using peroxidase-coupled anti-HA-Peroxidase antibodies (Roche, dilution 1/4000), peroxidase-coupled anti-GFP antibodies (Milteneyi, dilution 1/5000), anti-SUMO1 antibodies (69), anti-BES1 antibodies (11), anti-FBPase (Agrisera, dilution 1/5000). To quantify the ratio between BES1 and P-BES1, signal intensity obtained with anti-BES1 antibodies and corresponding to BES1 and P-BES1 was determined using Image J. Western blot analyses were performed in triplicates. Representative blots are shown in figures. For the loading control using anti-FBPase antibodies, the same membranes were stripped and used.

Immunoprecipitation experiments were carried out as previously described (25), using the μMACS GFP and HA isolation kits (Miltenyi Biotec). Input and immunoprecipitated fractions were separated by SDS-PAGE and subjected to western blot analyses as described above.

### Confocal microscopy

Dark-grown hypocotyls were treated with 100 µM CHX and 50 µM BFA for 15 min under vacuum before transfer to the corresponding temperatures prior to imaging. Hypocotyls were mounted in the same solution and imaged on a Leica TCS SP2 SP8 confocal laser scanning microscopes (www.leica-microsystems.com). The 514-nm laser line was used to image BRI1-mCit. Laser intensity and detection settings were kept constant in individual sets of experiments to allow the direct comparison of fluorescence levels. To image entire hypocotyl cells, the TileScan and Z-stack option was used. Due to the length of hypocotyl cells, the fluorescence intensity of the total plasma membrane (PM) was not possible. Rather, the intensity of different region of interest of equivalent size and corresponding to the plasma membrane was measured. The PM/BFA body ratio corresponds to the mean fluorescence of the PM portions and the mean fluorescence of BFA bodies. For localization and colocalization of transiently expressed proteins in *N. benthamiana*, the 488, 514, 561 nm laser line were used to image FLS2-GFP, BRI1-mCit and Desi3a-mCh, respectively. Colocalization analyses and determination of the Manders’ coefficient, highlighting the fraction of GFP/mCit signals colocalizing with mCh, were carried out using the ImageJ plugin JACoP (70).

### Statistical analyses

Data are shown as the average of three individual biological replicates, unless stated otherwise. Statistical analyses were performed with the software GraphPad Prism 7 software. Statistical significance of hypocotyl length between genotypes and/or conditions was assessed using one-way analysis of variance with post hoc Tukey test. Experiments had at least n=20 seedlings in each biological replicate. Quantification of western blots used the non-parametric Mann-Whitney (two genotypes/conditions) or Kruskal-Wallis (three genotypes/conditions and more) tests. Statistical significance is defined as follow: *, P ≤ 0.05; **, P ≤ 0.01; ***, P ≤ 0.001.

## Aknowledgements

We thank Cyril Zipfel and Yanhai Yin for sharing *bik1* mutant and anti-BES1 antibodies, respectively. We would also like to acknowledge the Imaging facility from the Fédération de Recherche Agrobiosciences Interactions et Biodiversité of Toulouse (FRAIB). This work was supported by research grants from the French National Research Agency (ANR-17-CE20-0026-01 to G.V.) and the French Laboratory of Excellence (project “TULIP” grant nos. ANR–10–LABX– 41 and ANR–11–IDEX–0002–02 to G.V.).

---

**Figure S1.**
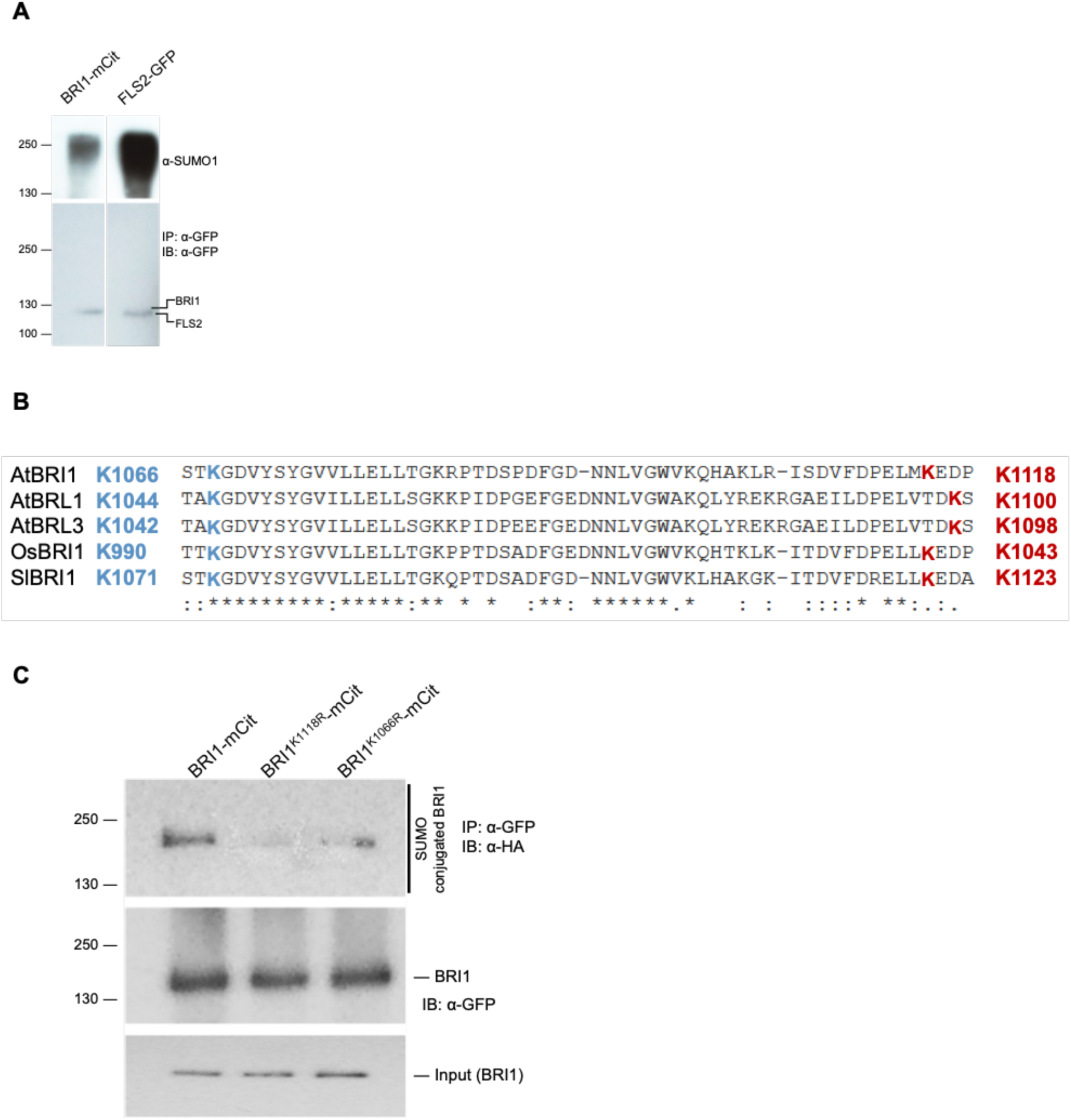
BRI1 is SUMOylated in vivo on two intracellular lysine residues. A. In vivo SUMOylation analyses of BRI1. Immunoprecipitation was carried out using anti-GFP antibodies on solubilized protein extracts from mono-insertional homozygous BRI1-mCitrine plants or the FLS2-GFP positive control plants. Detection of immunoprecipitated proteins used the anti-GFP (bottom), and anti-SUMO1 (top) antibodies. B. Sequence alignment of the cytosolic domain of BRI1 (amino acids 1064-1121) and its homologs BRL1 and BRL3. Predicted SUMO targets in BRI1 and homologs are shown. BC. In vivo SUMOylation analyses of BRI1, BRI1_K1066R_ and BRI1_K1118R_. BRI1-mCit, BRI1_K1066R_-mCit and BRI1_K1118R_-mCit were transiently expressed in *N. benthamiana* leaves prior to immunoprecipitation using anti-GFP antibodies on solubilized protein extracts. All constructs were co-expressed with SUMO1-HA. Detection of immunoprecipitated proteins used the anti-GFP (middle), anti-SUMO1 (top). The input fraction of BRI1 prior to immunoprecipitation is shown (bottom).

**Figure S2.**
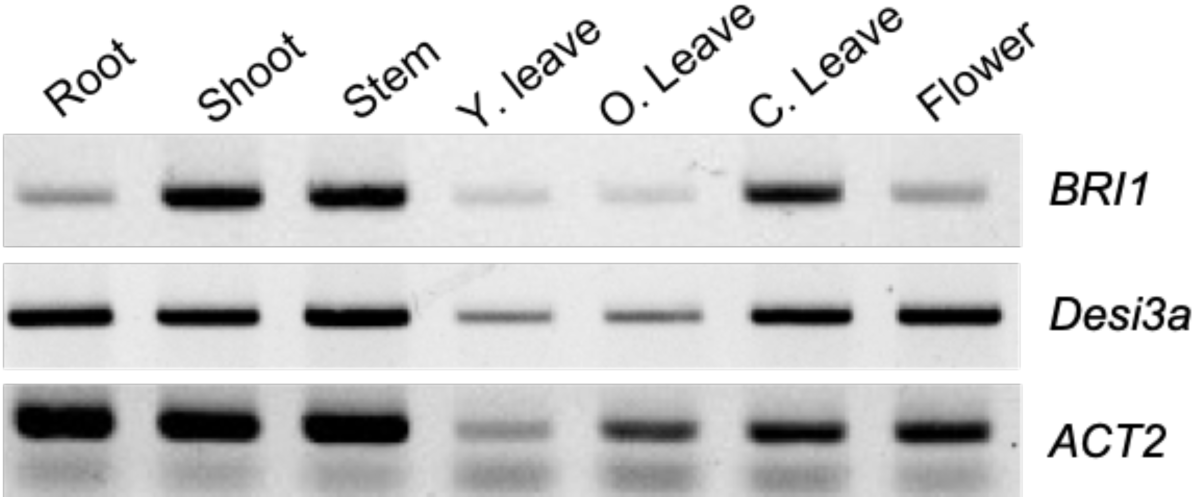
*Desi3a* and *BRI1* expression profiles overlap in plants. Semi-quantitative RT-PCR analyses of *Desi3a* and *BRI1* mRNA accumulation in different tissues of wild-type plants.

**Figure S3.**
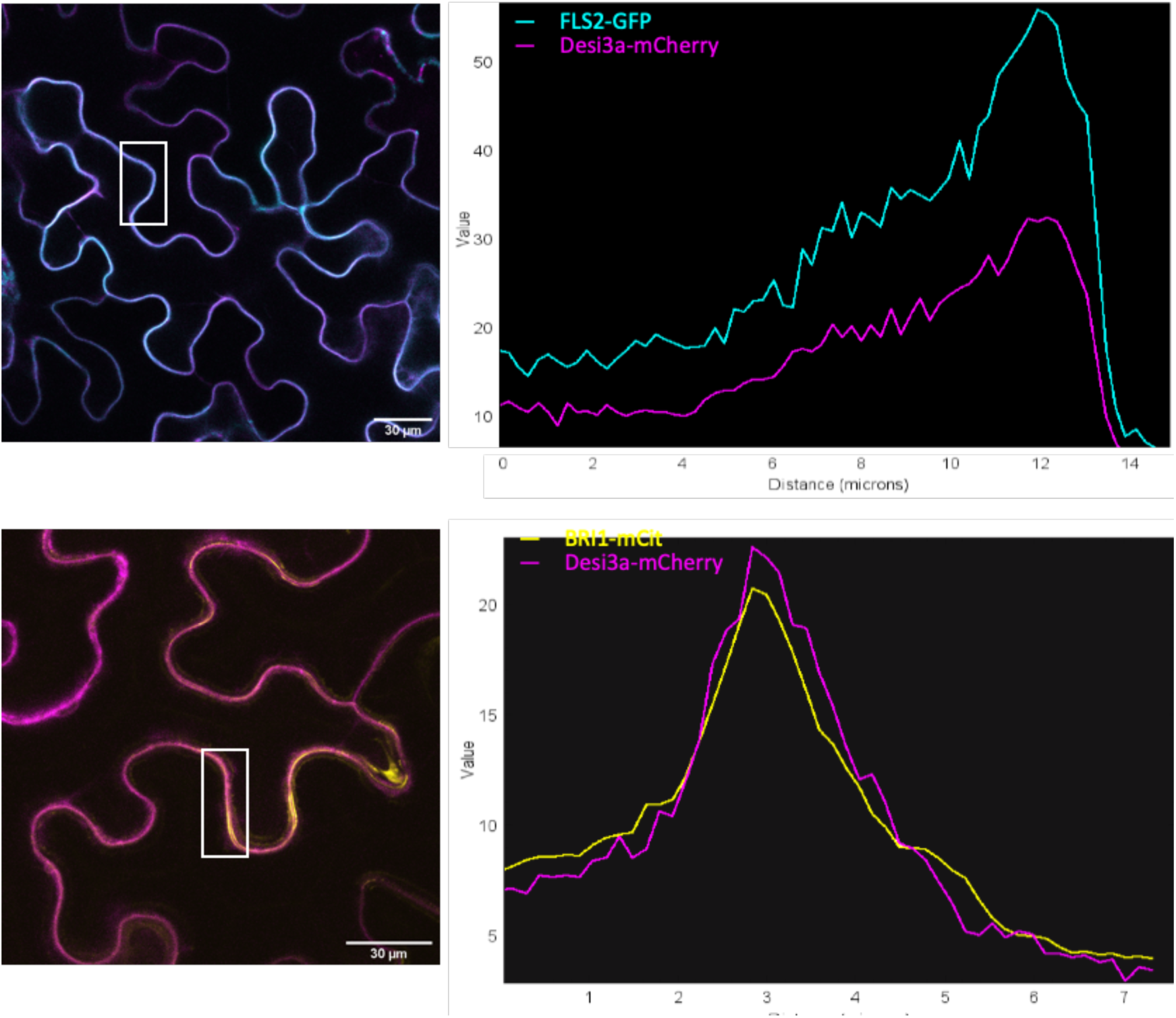
Colocalization of Desi3a with the plasma membrane-localized receptors FLS2 and BRI1. Fluorescence intensity profile of Desi3a-mCh with FLS2-GFP (top) and BRI1-mCit (bottom). The region of interest used to monitor fluorescence profiles are shown.

**Figure S4.**
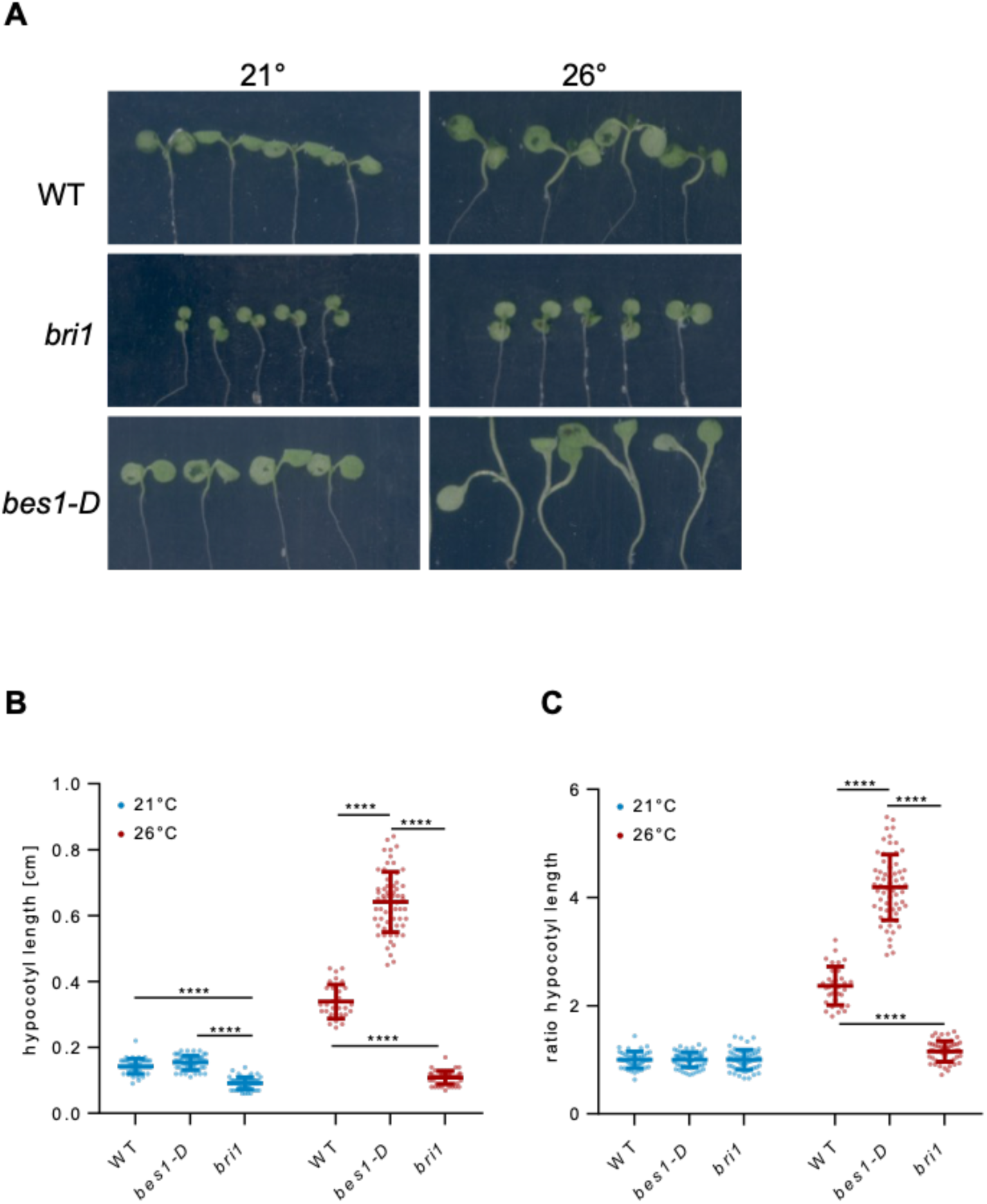
Phenotype of BR-related mutants at 21°C and 26°C. A. Hypocotyl length of 6-day-old wild-type (WT), *bri1* and *bes1-D* mutant plants grown at 21°C or 26 °C in the dark. Experiments were carried out in triplicates. Error bars represent SEM (n=20). The asterisk indicates a statistically significant difference with wild-type plants (two-way ANOVA with Sidak’s multiple comparison test). B. Ratio of hypocotyl length from wild-type (WT), *bri1* and *bes1-D* mutant plants grown at 21°C and 26 °C for 6 days. Experiments were carried out in triplicates. Error bars represent SEM (n=20). The asterisk indicates a statistically significant difference with wild-type (two-way ANOVA with Sidak’s multiple comparison test).

**Figure S5.**
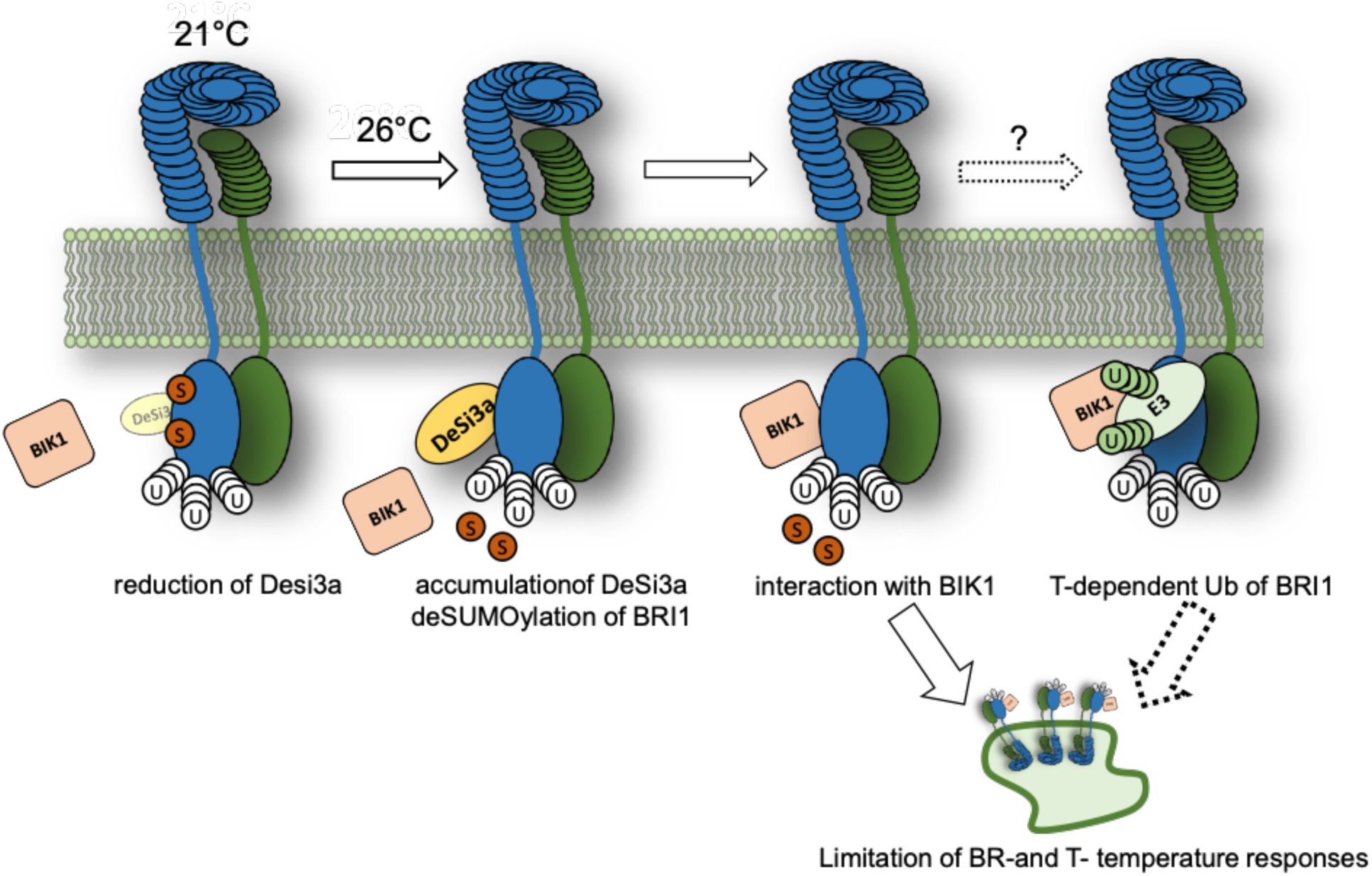
Model for the regulation by BRI1 by SUMO/deSUMOylation. Under standard temperature growth conditions (21°C), BRI1 is SUMOylated at residues K1066 and K1118 explained, at least in part, by the low levels of the Desi3a SUMO protease. Upon exposure to higher temperature (26°C), Desi3a accumulates and leads to the deSUMOylation of BRI1. DeSUMOylated BRI1 is downregulated through i) stronger interaction with the BIK1 negative regulator of BR signaling, and ii) increased ubiquitin-mediated endocytosis presumably through further ubiquitination of BRI1 at residues K1066 and K1118. This in turn limits the growth responses of hypocotyls to temperature elevation. Consequently, loss-of-function mutants for *desi3a* or *bik1* show enhanced responses to heightened temperature. BRI1 SUMO/deSUMOylation therefore acts as a temperature dependent switch that modulates BR-dependent responses and growth.

**Table S1.**
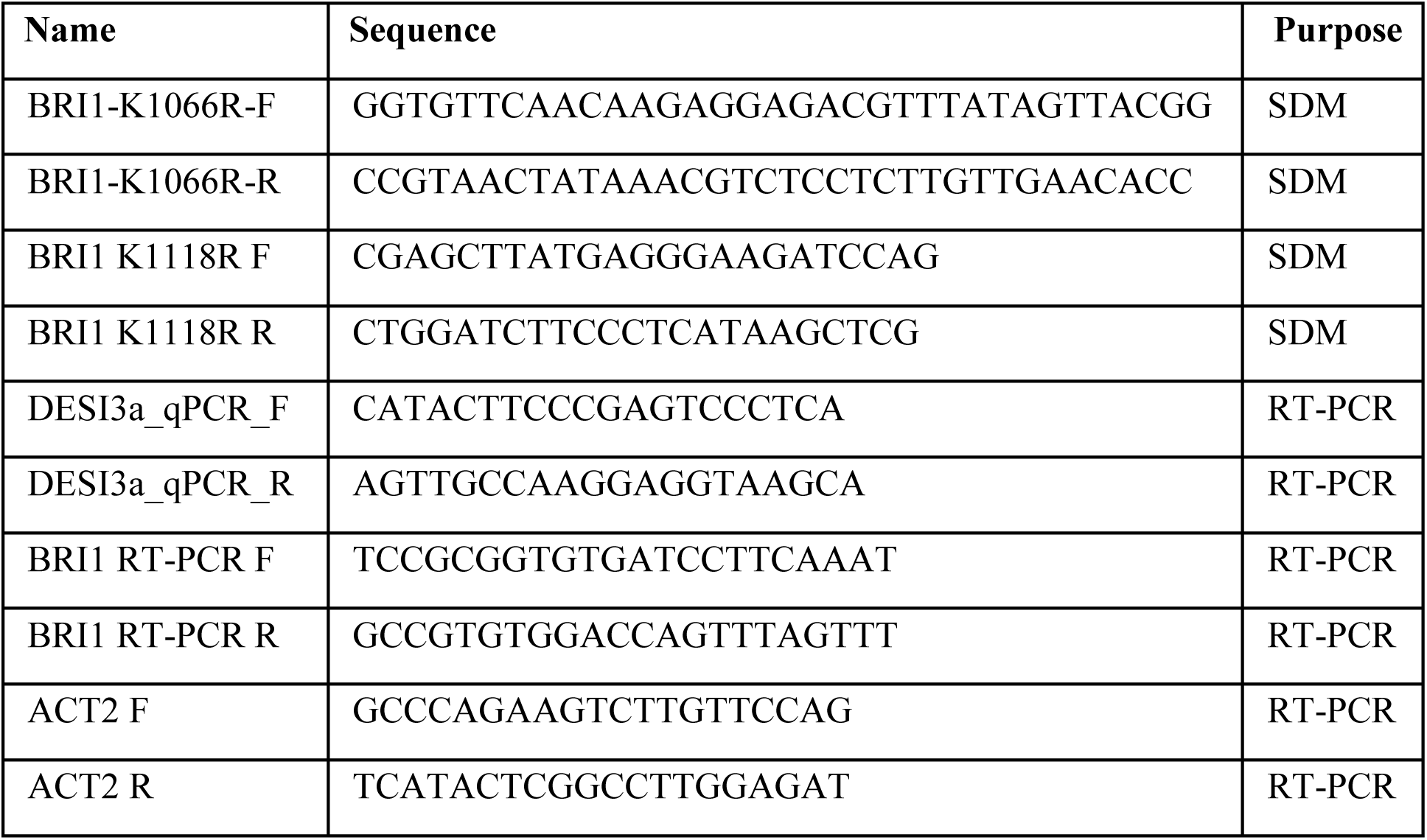
Primers used in this study

## References

1. Clouse SD (2011) Brassinosteroid signal transduction: from receptor kinase activation to transcriptional networks regulating plant development. Plant Cell 23(4):1219–1230.

2. He Z, et al. (2000) Perception of brassinosteroids by the extracellular domain of the receptor kinase BRI1. Science 288(5475):2360–2363.

3. Hothorn M, et al. (2011) Structural basis of steroid hormone perception by the receptor kinase BRI1. Nature 474(7352):467–471.

4. Kinoshita T, et al. (2005) Binding of brassinosteroids to the extracellular domain of plant receptor kinase BRI1. Nature 433(7022):167–171.

5. Li J & Chory J (1997) A putative leucine-rich repeat receptor kinase involved in brassinosteroid signal transduction. Cell 90(5):929–938.

6. She J, et al. (2011) Structural insight into brassinosteroid perception by BRI1. Nature 474(7352):472–476.

7. Wang ZY, Seto H, Fujioka S, Yoshida S, & Chory J (2001) BRI1 is a critical component of a plasma-membrane receptor for plant steroids. Nature 410(6826):380–383.

8. Li J, et al. (2002) BAK1, an Arabidopsis LRR receptor-like protein kinase, interacts with BRI1 and modulates brassinosteroid signaling. Cell 110(2):213–222.

9. Nam KH & Li J (2002) BRI1/BAK1, a receptor kinase pair mediating brassinosteroid signaling. Cell 110(2):203–212.

10. Wang ZY, et al. (2002) Nuclear-localized BZR1 mediates brassinosteroid-induced growth and feedback suppression of brassinosteroid biosynthesis. Dev Cell 2(4):505–513.

11. Yin Y, et al. (2002) BES1 accumulates in the nucleus in response to brassinosteroids to regulate gene expression and promote stem elongation. Cell 109(2):181–191.

12. Sun Y, et al. (2010) Integration of brassinosteroid signal transduction with the transcription network for plant growth regulation in Arabidopsis. Dev Cell 19(5):765–777.

13. Yu X, et al. (2011) A brassinosteroid transcriptional network revealed by genome-wide identification of BESI target genes in Arabidopsis thaliana. Plant J 65(4):634–646.

14. Clouse SD, Langford M, & McMorris TC (1996) A brassinosteroid-insensitive mutant in Arabidopsis thaliana exhibits multiple defects in growth and development. Plant Physiology 111(3):671–678.

15. Li J, Nagpal P, Vitart V, McMorris TC, & Chory J (1996) A role for brassinosteroids in light-dependent development of Arabidopsis. Science 272(5260):398–401.

16. Szekeres M, et al. (1996) Brassinosteroids rescue the deficiency of CYP90, a cytochrome P450, controlling cell elongation and de-etiolation in Arabidopsis. Cell 85(2):171–182.

17. Wang X, et al. (2005) Autoregulation and homodimerization are involved in the activation of the plant steroid receptor BRI1. Dev Cell 8(6):855–865.

18. Wang X & Chory J (2006) Brassinosteroids regulate dissociation of BKI1, a negative regulator of BRI1 signaling, from the plasma membrane. Science 313(5790):1118–1122.

19. Jaillais Y, et al. (2011) Tyrosine phosphorylation controls brassinosteroid receptor activation by triggering membrane release of its kinase inhibitor. Genes Dev 25(3):232–237.

20. Oh MH, Wang X, Clouse SD, & Huber SC (2012) Deactivation of the Arabidopsis BRASSINOSTEROID INSENSITIVE 1 (BRI1) receptor kinase by autophosphorylation within the glycine-rich loop. Proc Natl Acad Sci U S A 109(1):327–332.

21. Stone JM, Trotochaud AE, Walker JC, & Clark SE (1998) Control of meristem development by CLAVATA1 receptor kinase and kinase-associated protein phosphatase interactions. Plant Physiology 117(4):1217–1225.

22. Shah K, Russinova E, Gadella TW, Jr., Willemse J, & De Vries SC (2002) The Arabidopsis kinase-associated protein phosphatase controls internalization of the somatic embryogenesis receptor kinase 1. Genes Dev 16(13):1707–1720.

23. Wu G, et al. (2011) Methylation of a phosphatase specifies dephosphorylation and degradation of activated brassinosteroid receptors. Sci Signal 4(172):ra29.

24. Geldner N, Hyman DL, Wang X, Schumacher K, & Chory J (2007) Endosomal signaling of plant steroid receptor kinase BRI1. Genes Dev 21(13):1598–1602.

25. Martins S, et al. (2015) Internalization and vacuolar targeting of the brassinosteroid hormone receptor BRI1 are regulated by ubiquitination. Nat Commun 6:6151.

26. Zhou J, et al. (2018) Regulation of Arabidopsis brassinosteroid receptor BRI1 endocytosis and degradation by plant U-box PUB12/PUB13-mediated ubiquitination. Proc Natl Acad Sci U S A 115(8):E1906–E1915.

27. Martins S, et al. (2017) Brassinosteroid signaling-dependent root responses to prolonged elevated ambient temperature. Nat Commun 8(1):309.

28. Vierstra RD (2012) The expanding universe of ubiquitin and ubiquitin-like modifiers. Plant Physiol 160(1):2–14.

29. Srivastava AK, et al. (2018) SUMO Suppresses the Activity of the Jasmonic Acid Receptor CORONATINE INSENSITIVE1. Plant Cell 30(9):2099–2115.

30. Orosa B, et al. (2018) SUMO conjugation to the pattern recognition receptor FLS2 triggers intracellular signalling in plant innate immunity. Nat Commun 9(1):5185.

31. Tozluoglu M, Karaca E, Nussinov R, & Haliloglu T (2010) A mechanistic view of the role of E3 in sumoylation. PLoS Comput Biol 6(8).

32. Srivastava M, et al. (2020) SUMO conjugation to BZR1 enables Brassinosteroid signallint to integrate environmental cues to shape plant growth. Curr Biol:xxx-xxx.

33. Fei Q, et al. (2019) Effects of auxin and ethylene on root growth adaptation to different ambient temperatures in Arabidopsis. Plant Sci 281:159–172.

34. Rakusova H, et al. (2016) Termination of Shoot Gravitropic Responses by Auxin Feedback on PIN3 Polarity. Curr Biol 26(22):3026–3032.

35. Robinson DG, Jiang L, & Schumacher K (2008) The endosomal system of plants: charting new and familiar territories. Plant Physiol 147(4):1482–1492.

36. . Di Rubbo S, et al. (2013) The clathrin adaptor complex AP-2 mediates endocytosis of brassinosteroid insensitive1 in Arabidopsis. Plant Cell 25(8):2986–2997.

37. Lu D, et al. (2010) A receptor-like cytoplasmic kinase, BIK1, associates with a flagellin receptor complex to initiate plant innate immunity. Proc Natl Acad Sci U S A 107(1):496–501.

38. Lin W, et al. (2013) Inverse modulation of plant immune and brassinosteroid signaling pathways by the receptor-like cytoplasmic kinase BIK1. Proc Natl Acad Sci U S A 110(29):12114–12119.

39. Belkhadir Y & Jaillais Y (2015) The molecular circuitry of brassinosteroid signaling. New Phytol.

40. Singh M, Gupta A, & Laxmi A (2014) Glucose control of root growth direction in Arabidopsis thaliana. J Exp Bot 65(12):2981–2993.

41. de Lima CFF, Kleine-Vehn J, De Smet I, & Feraru E (2021) Getting to the Root of Belowground High Temperature Responses in Plants. J Exp Bot.

42. Fiorucci AS, et al. (2020) PHYTOCHROME INTERACTING FACTOR 7 is important for early responses to elevated temperature in Arabidopsis seedlings. New Phytol 226(1):50–58.

43. Gray WM, Ostin A, Sandberg G, Romano CP, & Estelle M (1998) High temperature promotes auxin-mediated hypocotyl elongation in Arabidopsis. Proc Natl Acad Sci U S A 95(12):7197–7202.

44. Jung JH, et al. (2016) Phytochromes function as thermosensors in Arabidopsis. Science.

45. Koini MA, et al. (2009) High temperature-mediated adaptations in plant architecture require the bHLH transcription factor PIF4. Curr Biol 19(5):408–413.

46. Legris M, et al. (2016) Phytochrome B integrates light and temperature signals in Arabidopsis. Science.

47. Stavang JA, et al. (2009) Hormonal regulation of temperature-induced growth in Arabidopsis. Plant J 60(4):589–601.

48. Sun J, Qi L, Li Y, Chu J, & Li C (2012) PIF4-mediated activation of YUCCA8 expression integrates temperature into the auxin pathway in regulating arabidopsis hypocotyl growth. PLoS Genet 8(3):e1002594.

49. Franklin KA, et al. (2011) Phytochrome-interacting factor 4 (PIF4) regulates auxin biosynthesis at high temperature. Proc Natl Acad Sci U S A 108(50):20231–20235.

50. Chung BYW, et al. (2020) An RNA thermoswitch regulates daytime growth in Arabidopsis. Nature plants 6(5):522–532.

51. Gonzalez-Garcia MP, et al. (2011) Brassinosteroids control meristem size by promoting cell cycle progression in Arabidopsis roots. Development 138(5):849–859.

52. Ibanez C, et al. (2018) Brassinosteroids Dominate Hormonal Regulation of Plant Thermomorphogenesis via BZR1. Curr Biol 28(2):303–310 e303.

53. Nemhauser JL, Mockler TC, & Chory J (2004) Interdependency of brassinosteroid and auxin signaling in Arabidopsis. PLoS Biol 2(9):E258.

54. Kruse M, et al. (2009) Impaired endocytosis of the ion channel TRPM4 is associated with human progressive familial heart block type I. J Clin Invest 119(9):2737–2744.

55. Cuijpers SAG, Willemstein E, & Vertegaal ACO (2017) Converging Small Ubiquitin-like Modifier (SUMO) and Ubiquitin Signaling: Improved Methodology Identifies Co-modified Target Proteins. Mol Cell Proteomics 16(12):2281–2295.

56. Chen J & Chen ZJ (2013) Regulation of NF-kappaB by ubiquitination. Curr Opin Immunol 25(1):4–12.

57. Desterro JM, Rodriguez MS, & Hay RT (1998) SUMO-1 modification of IkappaBalpha inhibits NF-kappaB activation. Mol Cell 2(2):233–239.

58. Irani NG, et al. (2012) Fluorescent castasterone reveals BRI1 signaling from the plasma membrane. Nat Chem Biol 8(6):583–589.

59. Robatzek S, Chinchilla D, & Boller T (2006) Ligand-induced endocytosis of the pattern recognition receptor FLS2 in Arabidopsis. Genes Dev 20(5):537–542.

60. Chinchilla D, et al. (2007) A flagellin-induced complex of the receptor FLS2 and BAK1 initiates plant defence. Nature 448(7152):497–500.

61. Heese A, et al. (2007) The receptor-like kinase SERK3/BAK1 is a central regulator of innate immunity in plants. Proc Natl Acad Sci U S A 104(29):12217–12222.

62. Sun Y, et al. (2013) Structural basis for flg22-induced activation of the Arabidopsis FLS2-BAK1 immune complex. Science 342(6158):624–628.

63. Schulze B, et al. (2010) Rapid heteromerization and phosphorylation of ligand-activated plant transmembrane receptors and their associated kinase BAK1. J Biol Chem 285(13):9444–9451.

64. Wang X, et al. (2008) Sequential transphosphorylation of the BRI1/BAK1 receptor kinase complex impacts early events in brassinosteroid signaling. Dev Cell 15(2):220–235.

65. Ortiz-Morea FA, He P, Shan L, & Russinova E (2020) It takes two to tango - molecular links between plant immunity and brassinosteroid signalling. J Cell Sci 133(22).

66. Zhang J, et al. (2010) Receptor-like cytoplasmic kinases integrate signaling from multiple plant immune receptors and are targeted by a Pseudomonas syringae effector. Cell Host Microbe 7(4):290–301.

67. Perraki A, et al. (2018) Phosphocode-dependent functional dichotomy of a common co-receptor in plant signalling. Nature 561(7722):248–252.

68. Karimi M, Depicker A, & Hilson P (2007) Recombinational cloning with plant gateway vectors. Plant Physiol 145(4):1144–1154.

69. Conti L, et al. (2008) Small ubiquitin-like modifier proteases OVERLY TOLERANT TO SALT1 and −2 regulate salt stress responses in Arabidopsis. Plant Cell 20(10):2894–2908.

70. Bolte S & Cordelieres FP (2006) A guided tour into subcellular colocalization analysis in light microscopy. Journal of microscopy 224(Pt 3):213–232.

